# Working and Reference Memory tasks trigger opposed long-term synaptic changes in the rat dentate gyrus

**DOI:** 10.1101/2020.08.02.230581

**Authors:** Mégane Missaire, Nicolas Fraize, Jean-Christophe Comte, Bruno Truchet, Régis Parmentier, Paul-Antoine Salin, Gaël Malleret

**Author notes:** corresponding author: Mégane Missaire and Gaël Malleret. **Email:**.

## Abstract

Long-term storage of information into memory is supposed to rely on long-term synaptic plasticity processes. Detection of such synaptic changes after training in long-term or reference memory (RM) tasks has yet been scarce, variable and only studied on a short time scale. On the other hand, short-term or working memory (WM) is largely known to depend on persistent neuronal activity or short-term plasticity processes. However, processing information into WM could also involve long-term synaptic changes that could be responsible for the erasure/forgetting of items previously stored in WM playing the role of proactive interference. In order to study long-term synaptic changes associated with RM or WM, we trained chronically implanted rats in three different radial maze tasks: a classical RM task and two WM tasks involving different levels of proactive interference. Synaptic responses at the perforant path to dentate gyrus synapse were recorded on a long-time scale (24h) in freely-moving rats after training in one of these three tasks. We found that consolidation of long-term information leads to a delayed synaptic potentiation, occurring 9 hours after RM training and predicting good behavioral performance on the following day. In contrast, optimal information processing into WM triggers a synaptic depression immediately observed after training and lasting 3 hours, that could act as a mechanism for interference erasure/forgetting.

## Introduction

Encoding and retrieving memories are fundamental processes for animal survival, especially when it comes to remembering the position of food sources in an environment. Such spatial memory processes mainly rely on the dorsal hippocampus, and have been suggested to depend on selective changes in synaptic transmission between related neurons (Takeuchi et al., 2014). Many studies linked such synaptic changes with memory processes using indirect, molecular or structural approaches (Fraize et al., 2017; Hayashi-Takagi et al., 2015; Matsuo et al., 2008; Ryan et al., 2015). However, a comprehensive, direct and unbiased account for the synaptic modifications occurring in real time in the brain after memory tasks is lacking. Moreover, while several types of memory have been described, it remains unclear whether they are supported by different synaptic plasticity processes.

In the taxonomy of memory, a major dichotomy has been established between long-term memory and short-term/working memory (WM). Long-term synaptic changes have mainly been studied and associated with long-term memory processes, based on the assumption that such processes should leave in the brain a durable trace (also called engram) (Tonegawa et al., 2015). It is commonly assumed that long-term memory relies on a sustained increase in synaptic transmission (similar to the one observed after an artificial induction of long-term potentiation or LTP (Bliss and Lømo, 1973)), and numerous attempts have been made to detect such synaptic changes in the brain after learning (Clarke et al., 2010; Fraize et al., 2017; Gruart et al., 2006; Pavlowsky et al., 2017; Takeuchi et al., 2014; Whitlock et al., 2006). The most direct method to detect such changes is certainly to record *in vivo*, in freely moving animals, evoked responses (field excitatory post-synaptic potentials, fEPSPs) at synapses of interest after training these animals in tasks involving long-term memory processes. However, the detection of these LTP-like changes after learning remains sparse and highly variable in amplitude (Gruart et al., 2015; Whitlock et al., 2006). Such differences could be explained by the time period during which fEPSPs were recorded. Indeed, fEPSPs were either recorded during the few minutes of the learning sessions (Gruart et al., 2006), or after memory tasks but only up to 4h (Whitlock et al., 2006) or 6h (Clarke et al., 2010) post-training. However, none of these studies recorded synaptic changes continuously beyond this time period. This lack of “long-term” studies is surprising as critical reorganizations related to memory consolidation have long been observed during specific time windows occurring up to 24h after learning (Smith, 1996).

Contrary to long-term memory, short-term or working memory (WM) is believed to rely not on long-term synaptic changes, but on the persistent activity of neurons encoding items stored during short delays of retention (Fuster and Alexander, 1971; Goldman-Rakic, 1995; Miyashita, 1988). However, this sustainable increase in firing rates may not be metabolically efficient and is not always observed during the whole retention delay (Rainer and Miller, 2002; Shafi et al., 2007). Therefore, some authors recently proposed a synaptic theory of WM (Mi et al., 2017; Mongillo et al., 2008), relying on a form of short-term presynaptic plasticity. However, no theory has ever suggested that long-term synaptic changes involving post-synaptic modifications could be associated with some aspects of WM. Indeed, WM is believed to be a short-term form of memory that once retrieved is supposed to be forgotten (Dudchenko, 2004), or transferred into long-term memory. Nevertheless, it is well known that WM is sensitive to proactive interference, such as past and irrelevant information previously stored in WM that can interfere with the recall of newer and more useful information (Roberts and Dale, 1981; Underwood, 1957; Wixted, 2004). Erasing – or inhibiting the recall of proactive interference would therefore be essential for WM function (Dudchenko, 2004). Such process could rely on long-term synaptic changes, especially those related to long-term synaptic depression (Fraize et al., 2017; Nicholls et al., 2008). However, until now, no study has ever followed the dynamics of long-term synaptic changes after WM training.

To palliate this lack of knowledge, we conducted a long-term electrophysiological study in freely-moving rats using an innovative comparative behavioral approach. Rats were trained in a radial maze in either a long-term – also called “*reference*” – memory (RM) task, or one of two WM tasks: a Low Interference (LIWM) or a High Interference WM task (HIWM) involving variable levels of proactive interference that differentially alter their level of performance (Fraize et al., 2016, 2017; Joseph et al., 2015; Missaire et al., 2017). Having previously shown that the dentate gyrus (DG) of the dorsal hippocampus (Fraize et al., 2017; Joseph et al., 2015) could play a major role in processing these interference, we performed long-term recordings of fEPSPs at the perforant path (PP) to DG synapse during two consecutive days. Our results show that while RM training promotes a delayed (>9h) increase in synaptic transmission that is positively correlated with next day performance of the trained rats, WM training also triggers long-term synaptic changes. We show that efficient processing of proactive interference in WM is correlated to a synaptic depression that is observed immediately after LIWM training and may support adaptive forgetting. In contrast, an inefficient processing of these interference during HIWM training depends on a delayed synaptic potentiation.

## Results

### A comparative approach to study reference memory, working memory and interference processing

The three behavioral tasks used in this experiment were performed in a 8-arm radial maze and involve the use of spatial, allocentric based (Bontempi et al., 1999; Maviel et al., 2004; Poirier et al., 2008) and hippocampus-dependent (Eichenbaum et al., 1999; O‟Keefe, 1993) memory. These tasks were designed in order to elicit the same level of motivation, locomotion of habituation in all trained animals. For instance, we controlled that the number of rewards collected and number of runs in the maze were equivalent for all three groups of rats and at any given session. Such controls allowed us to be certain that the difference observed at the synaptic level (evoked fEPSPs) between groups were directly linked to the cognitive process involved in each task (consolidation of long-term memory in RM *versus* processing of interference in WM).

Two of these tasks involved WM and were based on the same *delayed-non-match-to-place* (DNMTP) paradigm (see Materials and Methods) (Fig. 1*A*). However, the LIWM (Low Interference Working Memory) task was less repetitive than the HIWM (High Interference Working Memory) task. The pair of arms used in the LIWM task was thus different for each of the four trials of a given session. In contrast, during HIWM training, this pair of arms was always the same for all trials and sessions. It has consistently been found that the negative impact of proactive interference on WM increases with the level of similarity between items previously stored into WM (Cohen et al., 1994; Loess, 1968; Underwood, 1957). This is why the level of interference is considered higher in the HIWM task in which the same pair of arms is repeatedly presented to the rat. The third task used in this study was a classical RM (Reference Memory) task during which rats had to learn the fixed position of two food rewards in two arms of the maze (Fig. 1*A*).

**Figure 1.**
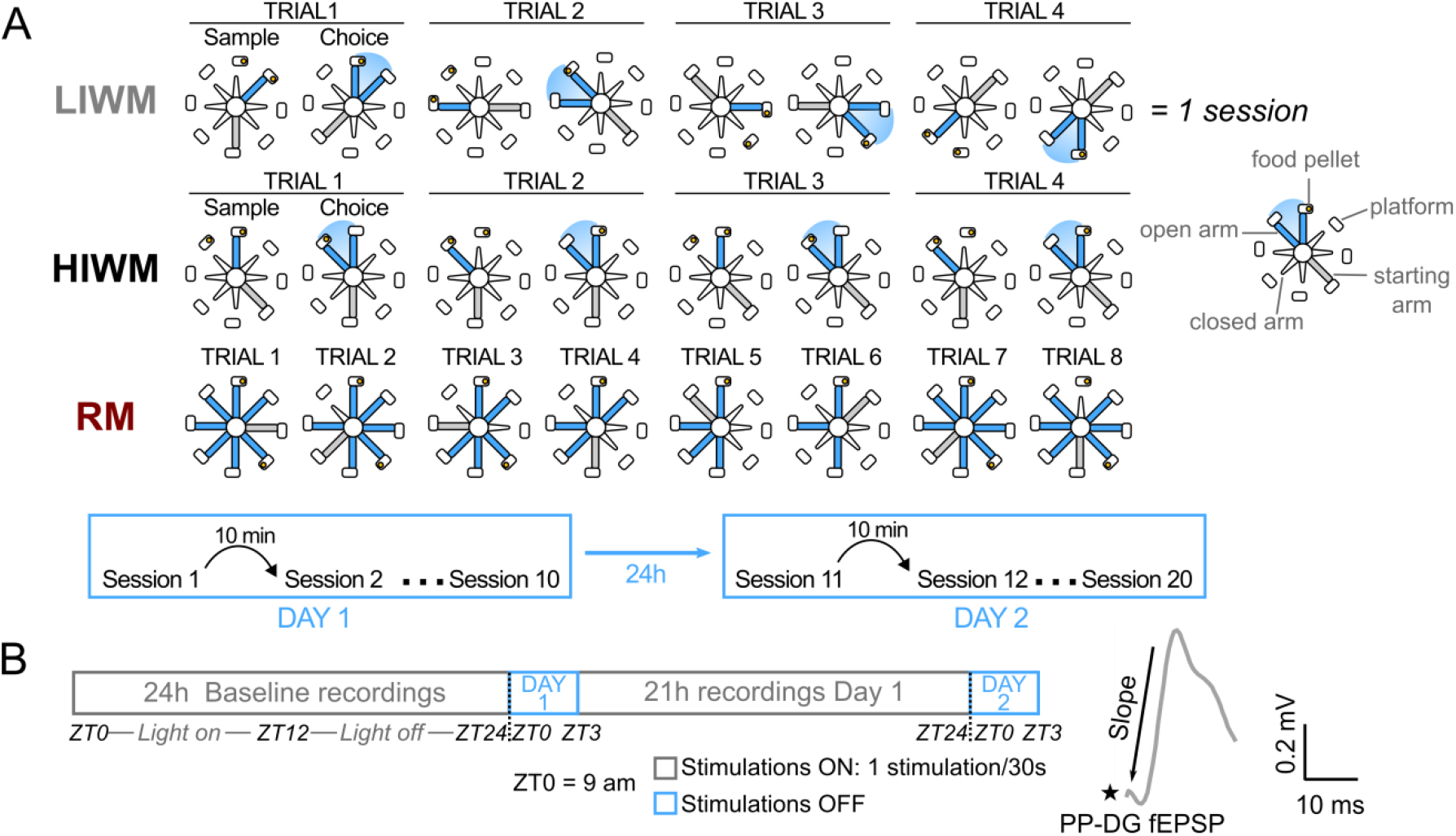
Behavioral tasks protocols and electrophysiological recordings. *(A)* Behavioral protocol for the three behavioral tasks: LIWM (Low Interference Working Memory), HIWM (High Interference Working Memory) and RM (Reference Memory) tasks – an example is given for one session in each task. The two WM tasks are based on a DNMTP paradigm with a sample and a choice phase, and the only difference between them is the use of different pairs of arms for each trial of a session in the LIWM task, contrary to the HIWM task during which the same pair of arms is used for all trials and all sessions. For the RM task, a possible sequence of runs chosen by an animal is represented, with the lowered arm on trial n being the one previously visited on trial n-1. In this example, the two rewards were collected on trial 6 and thus all arms were re-opened on trial 7. This task involves long-term storage and consolidation of a fixed rule (reference) in memory („*always go to the same two arms to get rewarded’*). The lower diagram depicts the training sequence common for the three tasks, with 10 sessions on Day 1 and 10 sessions on Day 2. *(B)* Recording protocol: fEPSPs at the PP-DG synapse were evoked every 30 seconds for 24h during the Baseline period (starting at ZT0 = 9am). Stimulations were then stopped during the 3 hours of training Day 1, and started again after the behavioral tasks for the remaining 21h (every 30s) before training Day 2 when stimulations were turned off again. The waveform is a classical fEPSP recorded at the PP-DG synapse, from which we computed the slope.

Chronically implanted rats (see Materials and Methods and next paragraph) were trained in these three tasks for 10 sessions per day on two consecutive days (Fig. 1*A*). Behavioral performance in these three tasks is expressed as the percentage of correct choices for blocks of two sessions, as displayed in Fig. 2*A*. For both WM tasks, performance was high (~70% of correct choices) at the very beginning of the task, probably because of the innate tendency of the rats to exhibit spontaneous alternation, a behavior that naturally causes rodents to choose a different option (visit arm #2) than the one previously adopted (visit arm #1), and in consequence to alternate exploration between two open arms (Tolman, 1925). For LIWM rats, performance increased across sessions on Day 1 (ANOVA 1: F_3.351,43.56_ = 7.406, p = 0.0003 ***), on Day 2 (ANOVA 1: F_2.183,28.38_ = 5.253, p = 0.0098 **), and also between Day 1 and Day 2 (Fig. 2*B*). On the contrary, for HIWM rats, no performance increase was observed during Day 1(ANOVA 1: F_2.910,40.75_ = 1.631, p = 0.1983), Day 2 (ANOVA 1: F_3.136,43.90_ = 1.615, p = 0.1980), or between Day and Day 2 (Fig. 2*B*). Consequently, the performance of HIWM rats was significantly lower than the performance of LIWM rats from the last block (B5) of Day 1 (p = 0.0235) and during Day 2. Such difference in the level of performance of rats trained in LIWM and HIWM tasks has systematically been observed in previous studies (Fraize et al., 2016, 2017; Joseph et al., 2015; Missaire et al., 2017) and likely reflects a non-optimal processing/forgetting of interference in rats training in the HIWM task. This inefficient processing of interference is further revealed by a between-session analysis of performance computed by trials instead of days (Fig. 2*C*). Such analysis shows that a difference in performance is observed between the two groups after a 24h-period of rest on the very first trials completed on Day 2 (Missaire et al., 2017). This result suggests that 24h-old information may impact the recall of recent (Day 2) information.

**Figure 2.**
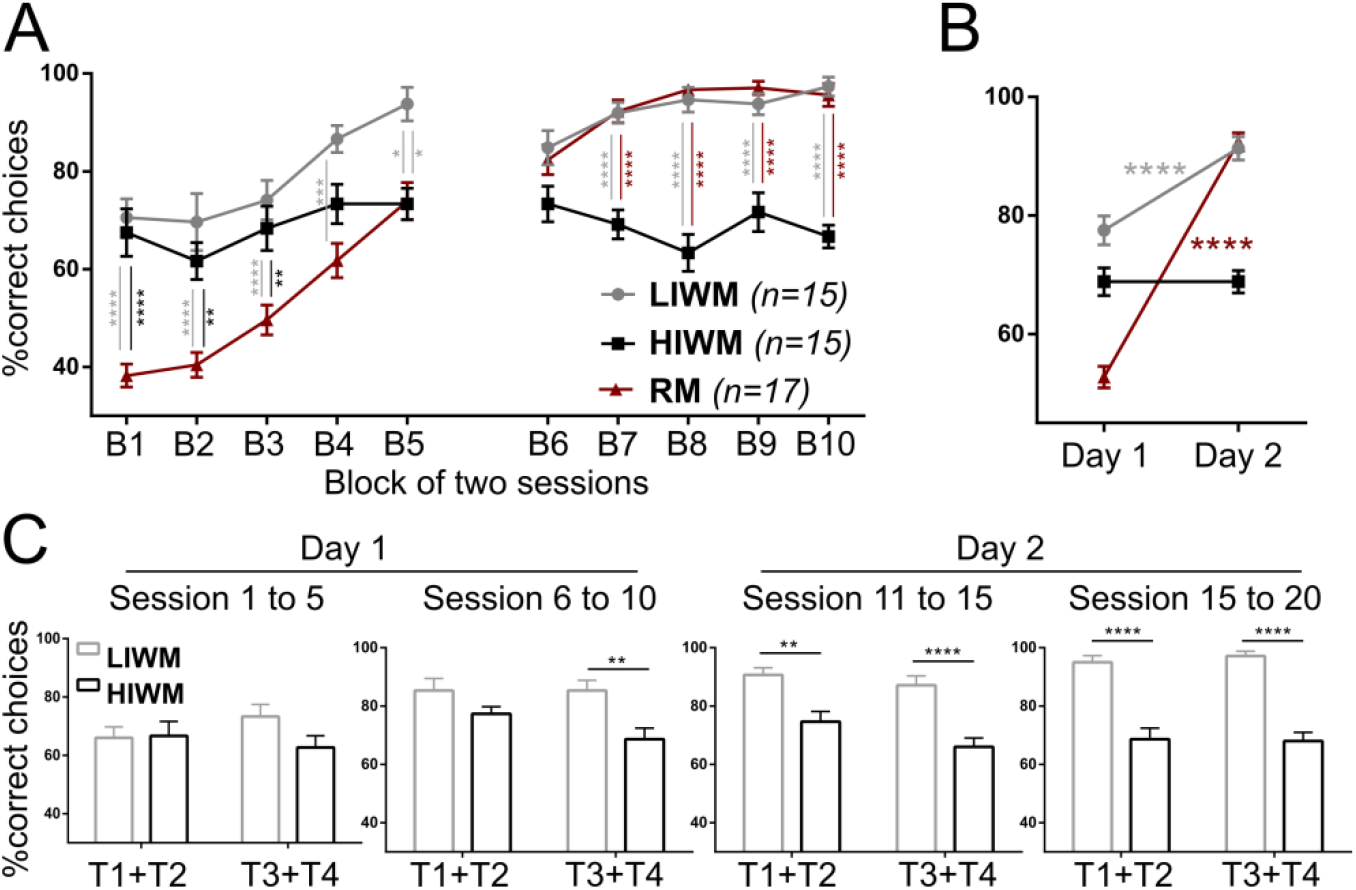
Three radial maze tasks involving working memory or long-term memory consolidation. *(A), (B)* Behavioral performance (percentage of correct choices) across the two days of training in the three tasks, averaged by blocks of two sessions *(A)* or by day *(B)*.Repeated-measure ANOVA revealed significant effects of Groups, Blocks/Day and Groups x Blocks/Day factors. More specifically, performance in the LIWM group is higher than in HIWM and improves between the two training days, while HIWM performance stagnates at a lower level across the two days of training. Performance in the RM group is lower than in the WM groups at the beginning of training, but rats rapidly learn the rule and improve their performance across the two days of training, reaching a level of performance equivalent to the one observed in LIWM rats on Day 2. *(C)* Within- and Between-session effect of proactive interference. Behavioral performance for the LIWM and HIWM groups averaged by two trials (two first or two last trials of a session) and by groups of 5 sessions. During the last 5 sessions of Day 1, performance of HIWM rats was lower than the one displayed by LIWM rats only for the last 2 trials (within-session effect), while on Day 2 this was the case from the very beginning (first trials) of the sessions (between-session effect). Data are represented as mean ± SEM.

In contrast, performance of rats trained in the RM task started at a lower level than the one observed in WM groups, below 40% and close to the chance level calculated at 34% for this RM task(Fraize et al., 2016). This task is indeed less „intuitive‟ for the rats and goes against their innate spontaneous alternation preference – see above. However, their performance rapidly increased during Day 1 (ANOVA 1: F_2.800,44.80_ = 27.14, p < 0.0001 ****), Day 2 (ANOVA 1: F_2.633,42.13_ = 12.82, p < 0.0001 ****), and between Day 1 and Day 2 (Fig. 2*B*).This increase in performance allowed RM rats to display on Day 2 a performance level equivalent to the one observed in LIWM rats (~100% at the end of Day 2) but higher than the one displayed by HIWM rats. To sum up, one group of rats succeeded in performing a WM task (LIWM), another failed to do so (HIWM), and a third group succeeded in learning a RM task.

We therefore used these behavioral groups to study and compare the long-term synaptic changes triggered by WM and RM tasks. We chose to record synaptic responses at the Perforant Path (PP) to Dentate Gyrus (DG) synapse, given the involvement of the DG in spatial long-term memory (Nanry et al., 1989; Xavier et al., 1999), spatial WM (Niewoehner et al., 2007; Sasaki et al., 2018; Xavier and Costa, 2009), and more specifically in the processing of proactive interference in spatial WM (Fraize et al., 2017; Joseph et al., 2015). Rats were therefore chronically implanted with a stimulating electrode in the PP and a recording array of 8 LFP electrodes in the DG (see Materials and methods). fEPSPs were continuously recorded before (24h Baseline period) and after (for 21h) training (that lasted 3h) on Day 1 (Fig. 1*B*).

### Opposed long-term synaptic changes after successful WM or RM training

Synaptic responses at the PP-DG synapse were obtained at the 8 electrodes implanted to target the DG for each rat. However, among these 8 electrodes, only recordings fulfilling precisely defined histological and electrophysiological criteria (see Materials and methods SI Appendix) were kept for analysis (corresponding to 6.7±1.4 standard deviation electrodes per rat in average). As shown in Fig. 3, the fEPSPs were relatively stable over the two days of recordings, indicating that the electrodes did not move despite the behavioral task.

**Figure 3.**
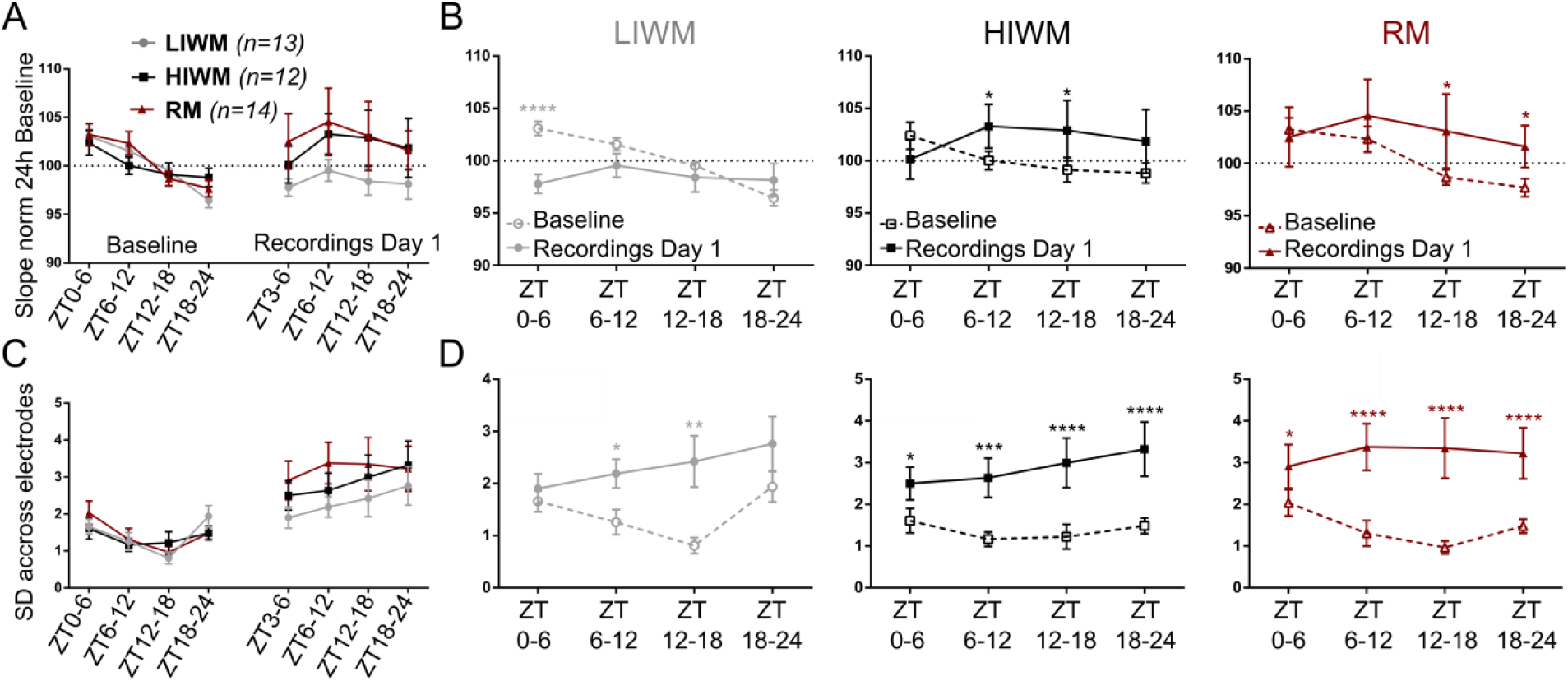
Working and reference memory tasks trigger opposed long-term synaptic changes with a local component. *(A)* fEPSP slope dynamics in the three groups during Baseline and Day 1 (post-training), averaged by 6h-periods and normalized to the averaged slope of the 24h of Baseline (dotted line). The repeated-measure ANOVA 2 did not reveal any Group of Time x Group significant effect. *(B)* Direct within-period comparison of the fEPSP slope (again normalized to the averaged slope on 24h of Baseline) between Baseline and the first day post-training. The repeated-measure ANOVA 2 and post-hoc tests revealed (i) a significant early synaptic depression after LIWM training (during the first 3h post-training – ZT3-6) compared to the same period in Baseline, (ii) a significant late synaptic potentiation in the HIWM group starting 3h and ending 15h post-training (ZT6-12 - ZT12-18), and (iii) a significant late synaptic potentiation in the RM group starting 9h and ending 21h post-training (ZT12-18 - ZT18-24). *(C)* Between-electrode and within-animal variability of the fEPSP slope during Baseline and Day 1. Evolution of the mean standard deviation of the fEPSP slope across electrodes of individual rats by 6-hours periods on Baseline and Day 1, for the three behavioral groups. No significant differences were observed between groups on Baseline or on Day 1. *(D)* For each of the three behavioral groups, comparison of the mean standard deviation between Baseline and Day 1 in each 6-hours period. This between-electrode and within animal variability of the fEPSP slope was significantly increased in all periods of Day 1 compared to Baseline in the three groups, except for the first period ZT0-6 in the LIWM group. Data are represented as mean ± SEM.

The slope of the PP-DG fEPSPs recorded in the three behavioral groups was averaged by 6-hour periods during Baseline and Day 1 (post-training) (Fig. 3*A*). The repeated-measure ANOVA 2did not reveal any significant effect of Group nor of the interaction Group x Time on Baseline or Day 1. However, a progressive decrease of the fEPSP slope could be observed in the three groups on Baseline. This decrease was not due to a progressive impairment of the synaptic responses, but to a circadian modulation of the synaptic responses at the PP-DG synapse (Fig. S1) that was previously described (Barnes et al., 1977). In order to take into account this circadian effect, we performed direct comparisons between the fEPSP slope recorded on Baseline and the one observed on Day 1 at the same time period, and we did so separately for the three behavioral groups (Fig. 3*B*). We thus observed an early and significant synaptic depression of 5% in the LIWM group on Day 1 (compared to Baseline), and more specifically in the first 3 hours following training (ZT3-6) (p < 0.0001). On the contrary, in the HIWM group, a significant synaptic potentiation of 3-4% was observed on Day 1 (compared to Baseline) much later on, between 3 and 15 hours post-training (ZT6-12: p = 0.0407, ZT12-18: p = 0.0142). Similarly, in the RM group, we observed a late synaptic potentiation on Day 1 (compared to Baseline), with an amplitude of 4%, occurring between 9 and 21h hours post-training (ZT12-18: p = 0.0198, ZT18-24: p = 0.0450). This analysis thus revealed opposed long-term modifications of the synaptic transmission at the PP-DG synapse after training in two WM tasks involving different levels of proactive interference (synaptic depression after LIWM training – synaptic potentiation after HIWM training). We can also note the opposed synaptic changes after successfully performing a WM task (LIWM – synaptic depression) or a RM task (synaptic potentiation).

These synaptic changes were monitored on average on all the reliable electrodes for a given rat. Interestingly, we also found that modifications of the synaptic transmission could be locally observed. Indeed, the standard deviation of the recorded fEPSP slope between-electrodes and within-animal was found to be significantly increased on Day 1 (compared to Baseline) at all time periods and in the three groups, except in the LIWM group for the first 3 hours immediately following training. Interestingly, this period was precisely the one during which we detected the synaptic depression in this group of rats (Fig. 3*C,D*). This result suggests that while training in general induces modifications at synapses more specifically engaged in the engram of a given memory (leaving other synapses unmodified), the synaptic depression after successful WM training (LIWM) is a more generalized process that can be recorded at multiple sites of recordings (at least in the DG).

### The long-term synaptic changes detected after WM or RM training are correlated with behavioral performance

We then assessed if the opposed long-term modifications of the synaptic transmission that we observed at the PP-DG synapse could be correlated with the behavioral performance of the rats. We first asked whether behavioral performance at the end of training on Day 1 could have an impact on the synaptic changes observed after training. We thus correlated the behavioral performance of the three groups of rats observed on Block 5 (last 2 sessions of Day 1) with the fEPSP slope in the following four 6-hour periods of Day 1 (normalized to 24h of Baseline fEPSP) (Fig. 4*A*). With each dot representing one animal on the illustrative scatter plot on the right, we could calculate Spearman’s rho, which was then plotted for each time period and behavioral group on the main graph on Fig. 4*A* *(left)*. A significant negative correlation (r = −0.334, p = 0.0186) was found between performance at B5 and fEPSP slope during the first 3 hours (ZT3-6) post-training in the LIWM group. In other words, LIWM rats with good performance at B5 display a more important synaptic depression during the first 3 hours post-training. For the HIWM group, we also observed a significant negative correlation (r = −0.573, p = 0.0264) between performance at B5 and fEPSP slope, but on later time period, at ZT12-18 on Day 1. As we showed that during this timeframe, HIWM rats display on average a synaptic potentiation (Fig. 3*B*), this late synaptic potentiation could be associated with bad performance on B5.

**Figure 4.**
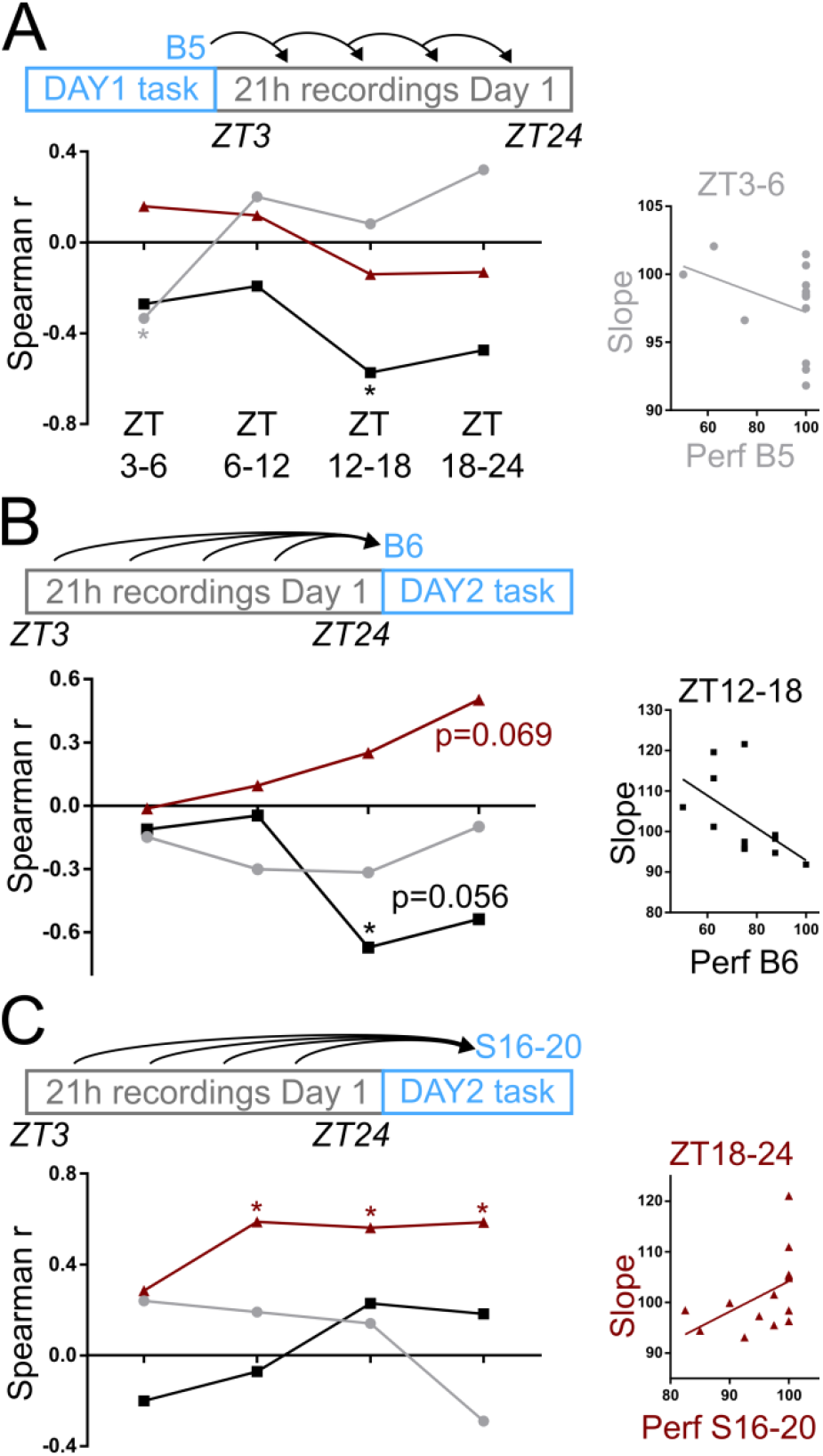
The recorded long-term synaptic changes are correlated with behavioral performance in the working and reference memory tasks. *(A)* Correlations between behavioral performance at the end of Day 1 (Block 5) and fEPSP slope recorded post-training during the four 6h-periods of Day 1 (normalized to the averaged fEPSP slope on 24h of Baseline). Note that good performance on B5 for LIWM rats predict synaptic depression during the first 3h post-training (negative correlation between past performance and fEPSP slope: r = −0.334, p = 0.0186 *), as illustrated by the scatter plot on the right (each dot representing one animal). *(B)* Correlations between fEPSP slope recorded post-training during the four 6h-periods of Day 1 (normalized to the averaged fEPSP slope on 24h of Baseline) and behavioral performance at the beginning of Day 2 (Block 6). Note that synaptic potentiation at the end of post-training Day 1 (more specifically at ZT12-18) for HIWM rats predicts bad behavioral performance on B6 (negative correlation between fEPSP slope and future performance: r = −0.672, p = 0.0128 *), as illustrated by the scatter plot on the right (each dot representing one animal). *(C)* Correlations between fEPSP slope recorded post-training during the four 6h-periods of Day 1 (normalized to the averaged fEPSP slope on 24h of Baseline) and behavioral performance at the end of Day 2 (Sessions 16-20). Note that synaptic potentiation from 3 to 21h post-training Day 1 in the RM group predicts good behavioral performance on S16-20 (positive correlations between fEPSP slope and future performance for ZT6-12: r = 0.5875, p = 0.0301 *; ZT12-18: r = 0.5616, p = 0.0396 *; ZT18-24: r = 0.581, p = 0.0309 *), as illustrated by the scatter plot on the right for ZT18-24 (each dot representing one animal). Data are represented as mean ± SEM.

We also asked whether synaptic changes occurring after training on Day 1 predict behavioral performance on Day 2.First, we correlated the fEPSP slope in the four 6-hour periods of Day 1 (normalized to 24h of Baseline fEPSP) with behavioral performance at Block 6 (two first sessions of Day 2) for the three behavioral groups (Fig. 4*B*). For the HIWM group, a significant negative correlation was found between the fEPSP slope at ZT12-18 of Day 1 (r = −0.672, p = 0.0128) and performance at B6. Note that the correlation tends to extend to ZT18-24 (r = −0.5382, p = 0.0562). Therefore, the synaptic potentiation we observed at ZT12-18 for the HIWM group appears to be correlated with decreased performance on Day 2. On the contrary, for the RM group, even if not significant, a positive correlation between fEPSP slope in the last 6 hours of Day 1 (ZT18-24) and performance at B6 (r = 0.5023, p = 0.0693) can be observed. We thus wanted to know if this possible link between potentiation and superior performance in RM could be stronger at the end of training Day 2, when rats typically reach a plateau and a maximum of performance. We thus performed the same correlations, but this time with average performance during the last 5 sessions of Day 2 (S16-20) (Fig. 4*C*). With this new analysis, we observed significant positive correlations between fEPSP slope at ZT6-12 (r = 0.5875, p = 0.0301), ZT12-18 (r = 0.5616, p = 0.0396), ZT18-24 (r = 0.5851, p = 0.0309) and performance at S16-20 for the RM group. The synaptic potentiation observed during these time periods on Day 1 (Fig. 3*B*) could thus be associated with a stronger memory consolidation of the rule to be learned in the task.

## Discussion

Our findings show that synaptic changes at the PP-DG synapse can be detected after both spatial RM and WM training. These changes are 1) long-lasting – they are observed for at least 3h and can last more than 6h; 2) often delayed in time – they occur 6 to 15h after training in some cases; and 3) bidirectional – they can lead to a synaptic depression or synaptic potentiation depending on the cognitive process involved.

### Synaptic depression – An adaptive forgetting marker?

We show that WM training can induce long-term changes in synaptic transmission. Synaptic transmission can thus be decreased or increased whether training (LIWM or HIWM) involved a more or less important processing of proactive interference. More specifically, LIWM training that is associated with successful processing of interference (Fig. 2), is followed by, and correlated with, a synaptic depression (Fig. 3 and Fig. 4*A*). As we assessed the early slope of fEPSPs, it is unlikely that this synaptic depression could be the result of a long-lasting increase in GABAergic inhibition. On the contrary, this depression would depend on a molecular cascade associated with LTD-like processes as suggested by a recent study from our team (Fraize et al., 2017). This western blot showed that LIWM training is associated with a significant decrease in the phosphorylation of the α-amino-3-hydroxy-5-methyl-4-isoxazolepropionic acid receptors (AMPARs) GluA1 subunits at the Ser-845 site. Ser-845 GluA1 dephosphorylation is considered to be a signal for AMPARs internalization and their lysosomal degradation, that *de facto* decreases synaptic transmission during LTD (Fernández-Monreal et al., 2012). Since our present *in vivo* findings show that synaptic depression is immediately observed after LIWM training (ZT3-6), we believe this depression reflects the late phase of an LTD induced in the DG while the rats were still performing the task. This online process (during behavioral training) could reflect adaptive forgetting of prior irrelevant information (pertaining to past trials) and a “reset” of WM content (Roberts and Dale, 1981) so that new information may be stored once more in WM during subsequent trials. Rats having the best performance on Day 1 would be those with the most effective adaptive forgetting of WM content, and therefore those displaying the highest LTD (as suggested by our correlation results - Fig. 4*A*).

This interpretation is in agreement with previous work suggesting that LTD would be involved in forgetting processes, and more specifically adaptive forgetting. We previously showed that the genetic inhibition of the Protein Phosphatase 2A (PP2A) that blocks LTD and AMPARs internalization leads to the inhibition of adaptive forgetting of past information in both a flexibility-based RM water maze task or a WM T-maze task (Nicholls et al., 2008). In contrast, inhibiting phosphorylation processes by genetically expressing a dominant negative form of the regulatory subunit of Protein Kinase A (PKA) enhances LTD, flexible learning and the adaptive forgetting of interference in a radial maze WM task similar to the one used in the present study (Malleret et al., 2010). Recently, Malinow and colleagues (Nabavi et al., 2014) showed that the optogenetic delivery of an LTD conditioning protocol targeting the auditory input in the amygdala leads to the forgetting of the association between a foot shock and this auditory input previously learned by a rat. Others found that blocking AMPARs internalization prevents normal forgetting of established long-term memories for object location (Migues et al., 2016) or the flexible learning of new water maze locations in RM or WM due to an absence of forgetting of previously learned past positions (Awasthi et al., 2018). Altogether, these results could explain why the efficient processing (forgetting) of interference may prompt LTD-like phenomenon that is observed immediately after LIWM training.

Although prolonged (lasting around 3h), this synaptic depression may be considered of a rather weak in amplitude (a fEPSP slope decrease of 5% compared to baseline). However, even if the group of Manahan-Vaughan studied extensively the artificial induction of LTD in the hippocampus and its modulation after various learning tasks (Kemp and Manahan-Vaughan, 2007), no physiological LTD-like process has ever been clearly reported *in vivo* after behavioral training. Consequently, it is difficult to compare our findings with others from the scientific literature. Even the artificial induction of homosynaptic LTD at the PP-DG synapse seems difficult to obtain *in vivo*. Indeed, a heterosynaptic LTD can be easily induced in the medial perforant pathway (MPP) (by inducing LTP on the lateral perforant pathway (LPP)), with an amplitude of 30% of synaptic depression (measured on the slope of the fEPSPs) and lasting at least 3 days (Doyère et al., 1997). However, the direct induction of homosynaptic LTD in one of these pathways leads to a 5 to 10% synaptic depression in some (but not all) animals, lasting only 60 minutes approximately (Gonzalez et al., 2014). The strength or the present study is that the synaptic depression observed is not induced artificially but physiologically induced, after training in a task that naturally involves adaptive forgetting. As this forgetting process would allow previously stored information (related to past trials) to be put aside or even erased, we can assume that the LTD-like phenomenon we observe corresponds more to a depotentiation rather than a *de novo* LTD, that would still share with LTD common mechanisms but that is easier to induce than LTD *in vivo* (Bear and Abraham, 1996). Indeed, some authors have suggested that storage of information in WM would imply synaptic potentiation, and more specifically short-term potentiation (STP – (Park et al., 2014)). Depotentiation would be the mechanism by which this stored information would be forgotten/erased.

Is this synaptic depression a local or widespread mechanism? To test this hypothesis, we computed the average intra-animal and inter-electrode standard deviation to assess differences between electrodes (and therefore synapses) in the DG. We then compared this standard deviation between Baseline and Day 1 by 6h time slots (Fig. 3*D*). Interestingly, we observe that this variability between recording electrodes is not increased (as it is the case in the other groups or time slots) following LIWM training compared to Baseline in the first 3h period, that corresponds to the time slot during which the synaptic depression is observed. This result suggests that this synaptic depression could thus be generalized to multiple DG synapses rather than being a local phenomenon. One possible interpretation for this result is that repeating numerous trials (40 per day) during LIWM training causes massive storage of information in WM (involving numerous synapses), that would subsequently require massive adaptive forgetting (at the level of the same important number of synapses) to allow deletion (and “reset”) of this information in WM, potentially through a generalized and prolonged LTD-like process like the one we observe.

In contrast, the variability between recording electrodes is increased after HIWM training (compared to Baseline) for all time periods of post-training recording (Fig. 3*D*). This suggests the existence of localized, rather than generalized, modifications in the synaptic transmission in this task. Contrary to what is observed after LIWM training, synaptic transmission is potentiated (by approximately 4%) after HIWM training, and in particular 3 to 15h post-training (Fig. 3*B*). This potentiation is in agreement with previous results from our team (Fraize et al., 2017) showing that HIWM training increases GluA1 expression in the DG, as well as the level of phosphorylation of this AMPAR subunit. A massive increase (+ 60%) in the phosphorylation of Ca2+/calmodulin-dependent protein kinase II (CaMKII) in the DG was also observed in the same study. It has been shown that phosphorylation of CaMKII constitutes an important signal for the induction of LTP, allowing phosphorylation and exocytosis of GluA1-AMPARs at the synaptic level, and *de facto* the increase/potentiation of synaptic transmission (Citri and Malenka, 2008). As an increase in GluA1 expression is also observed after HIWM training (Fraize et al., 2017), we can hypothesize that this task promotes the establishment of a late phase of LTP that would require the synthesis of new protein, and in particular new AMPARs. The present *in vivo* observation of a prolonged (long-term) increase in synaptic transmission after HIWM training seems to confirm the existence of LTP-like phenomena after completion of this task.

This synaptic potentiation is also negatively correlated with performance on Day 2 (Fig. 4*B*), suggesting that this potentiation is deleterious for behavioral performance, and thus for adaptive forgetting. Unlike the synaptic depression observed after LIWM training allowing previously stored information (Day 1) to be put aside or erased, this LTP-like phenomenon after HIWM training could prevent such forgetting process, and contribute to the consolidation of this past information (Day 1) playing the role of interference on the next day (Day 2). This is suggested by the analysis of the behavioral performance of rats trained in the HIWM task. While performance of HIWM rats is lower than the one displayed by LIWM rats only for the last 2 trials (within-session effect) of Day 1, this is the case on Day 2 from the very beginning (first trials) of the training sessions (between-session effect)(Fig. 2*C*). This result suggests a lack of WM reset for rats trained in the HIWM task: they display impaired performance as soon as training starts on Day 2 because past information (Day 1) has not been put aside/erased from WM (reset) and has even possibly been consolidated in long-term memory (Missaire et al., 2017; Roberts and Dale, 1981). This consolidation of past information (playing the role of interference on Day 2) would be achieved through a delayed synaptic potentiation (occurring several hours after the end of training) that would have a deleterious effect on the performance of the rats trained in the HIWM task.

### Consolidation of information into long-term memory (RM) and delayed LTP

In contrast to the HIWM group, the synaptic potentiation observed after RM training (Fig. 3*B*) has a positive effect on the behavioral performance of the rats trained in this task the next day (positive correlation between the synaptic responses recorded on Day 1 and the rats performance in RM on Day 2 - Fig. 4*C*). Indeed, RM and HIWM training are quite different and even opposite in nature. While the consolidation of past information (Day 1) interferes with the recall of newer information (Day 2) in the HIWM task, this consolidation of past information is required for acquiring the rule (invariant information) during incremental learning of the RM task. This interpretation is in agreement with several *in vivo* studies showing that training involving long-term memory formation increases synaptic transmission (Clarke et al., 2010; Gruart et al., 2006; Moser et al., 1994; Whitlock et al., 2006). However, most of these studies recorded fEPSPs during the first hours following training, or even for some, only during training itself. The originality of our approach was to continuously record synaptic changes after the first day of RM training. This is how we were able to show that synaptic responses can be durably increased (LTP-like phenomenon) several hours after training (delayed-LTP). If we had only recorded fEPSPs during the first hours post-RM training, we would not have detected any change in synaptic transmission. This could explain why, while there is considerable evidence in the literature that altering molecular mechanisms associated with LTP can prevent memory processes, there is in fact relatively few evidences of LTP-like phenomenon after behavioral training. Consequently, the difficulty to find such alterations in synaptic transmission after training in tasks involving long-term memory could be partly due to the time window analyzed in these studies. This synaptic potentiation delay may seem odd and could lead to reconsider the contribution of DG and/or LTP in RM. However, a lesion of the DG shows that this structure is required in RM (Joseph et al., 2015). Furthermore, as pointed out above, there is a positive correlation this delayed synaptic potentiation and performance at D2. It is known that critical reorganizations of sleep, known to play an important role in memory consolidation (Rasch and Born, 2013) as well as in synaptic plasticity (Diekelmann and Born, 2010; Tononi and Cirelli, 2014), can take place up to 24 hours after RM training (Smith, 1996). Consequently, the same delayed reorganizations in synaptic transmission could occur after training. Our study brought to light such delayed synaptic modifications following training in a long-term RM task. However, we cannot exclude the fact that an earlier synaptic potentiation takes place but is not recorded at the level of our electrodes, that *in fine* only correspond to a small portion of the rat‟s DG.

In our study, the synaptic potentiation observed after RM training is rather weak, representing an increase in the fEPSP slope of 4% compared to baseline. However, this weak potentiation is certainly much more compatible with physiological processes of memory storage than the massive potentiation (sometimes above 100%) observed *in vivo* after artificial inductions of LTP. In addition, to detect these synaptic changes, we used extracellular recordings. fEPSPs correspond to the summation of extracellular synaptic currents of multiple synapses localized in the vicinity of the recording electrode (Chaillan et al., 2008). Therefore, large increase in synaptic transmission to individual synapses could be partially balanced at the population level by null or opposed modifications at other synapses. Engram cells studies suggest that only 10 to 30% of the neurons of a given circuit would be recruited for the storage of a given memory. For instance, 20% only of hippocampal CA1 pyramidal cells express the immediate early gene *c-Fos* after exploration of a new environment (Tanaka et al., 2018). Furthermore, only a fraction of these engram cells‟ synapses would be modified by memory. Matsuo and colleagues have shown that the increase in synaptic GluA1 recruitment in CA1 after contextual fear conditioning barely affects 3% of dendritic spines (Matsuo et al., 2008). Similarly, after training in an inhibitory avoidance task, Bear and colleagues detected synaptic potentiation in CA1 in only 12 out of 44 recording electrodes, other synapses remaining stable or becoming durably depressed (Whitlock et al., 2006).

Our findings are in line with these results. Not only RM training induces a rather weak (but still prolonged) synaptic potentiation (4% compared to baseline), but this potentiation is delayed in time, extremely variable (as indicated by the increase in the variability between recording electrodes - Fig. 3*D*) and can be reversed by subsequent RM training. Indeed we found that the late synaptic potentiation observed at the end of Day 1 can be replaced by a synaptic depression (compared to Baseline and compared to Day 1) after training Day 2 (Fig. S2). This result is in agreement with previous results from our team showing that overtraining rats in a spaced (1 session per day) RM task induces a decrease (and not an increase as expected) in the expression of LTP markers (phosphorylation of CaMKII) in the DG (Fraize et al., 2017). The synaptic depression we observe after 2 days of RM training (Fig. S2) could participate in the abstraction/semantization of the memory acquired during the RM task. In fact, during RM training, rats would have first to extract and store (through synaptic potentiation?) a general rule of the task by processing invariant information during multiple episodes or trials (“*food is located in arms # 1 and # 4 of the maze*”). Later on however, they would also have to forget (through synaptic depression?) non-essential information (related for instance to the internal state of the animal during a specific trial) in order 1) not to overflow their brain with irrelevant information and 2) keep a flexible use of the general rule previously acquired. This is also how semantic memory would be generated out of multiple experiences originally processed in episodic memory (Hardt et al., 2013). This episodic-to-semantic shift would be very similar to context generalization, the tendency to express a behavior that was once specific to a specific learning context in other contexts. Hardt and colleagues (Migues et al., 2016) showed that blocking AMPARs internalization not only prevents normal forgetting of established long-term memories, but also the generalization of contextual fear expression (the natural tendency for an animal to express after several days fear for other contexts than the one originally trained for). Consequently, this result as well as ours suggest that LTD mechanisms could participate to the formation and transformation of RM, from an episodic-based to a semantic-based memory trace.

## Materials and Methods

### Electrode implantation for chronic recordings

Stereotaxic surgery was performed on 33 Dark Agouti male rats aged 10 weeks. Surgical procedures are further described in detail in SI Appendix. A recording array of 8 LFP electrodes was implanted in the molecular layer of the DG, and a bipolar stimulating electrode was implanted in the PP, coming from the entorhinal cortex. In addition, two EEG electrodes were fixed on the skull (prefrontal and parietal), two EMG electrodes were inserted between the neck muscles, and a reference electrode was fixed above the cerebellum for referential recordings.

### Recording setup and electrical stimulations

The detail of the recording setup is described in SI Appendix, and rats remained connected to the setup during the whole duration of an experimental protocol. Electrical stimulations consisted in a monophasic 200μs pulse delivered by an isolated pulse stimulator (model 2100, *AM-Systems, U.S.A*.). Before all experimental protocols, an input-ouput (I/O) curve was established in order to choose the stimulation intensity used for the whole experiment. This intensity had to elicit an fEPSP with an amplitude corresponding to 75% of the maximal amplitude on the 8 electrodes. During the experimental protocol, stimulations were elicited every 30 seconds, except during the behavioral tasks when stimulations were stopped. Continuous recordings of stable fEPSPs were acquired for 24h before the beginning of the behavioral task and were used as a baseline.

### Behavioral tasks and protocol

The three behavioral tasks have been used in previous studies (Fraize et al., 2016, 2017; Joseph et al., 2015), and are described in more details in SI Appendix. Briefly, these three appetitive spatial tasks took place in the same radial maze and were equivalent in terms of duration, locomotion, food rewards, or motivation. The two WM tasks were based on a DNMTP (Delayed Non-Match to Place) paradigm, with 1) a sample phase during which the animal was forced to visit the only open arm of the maze containing a first food reward, and 2) a choice phase during which the animal had to choose between entering the arm previously visited during the sample phase (familiar arm – wrong choice) that no longer contained any reward, or entering an adjacent new arm (correct choice) containing a second food pellet. These two phases represented one trial, and were separated by a 15 second-delay (during which rats were placed in an opaque transfer cage on the central platform of the maze), compatible with the storage duration of information (here, the arm visited during the sample phase) in WM. After the end of a given trial, this piece of information stored in WM is no longer useful and needs to be forgotten in order not to interfere proactively with the storage and recall of newer information in WM during the next trials. Indeed, four trials were performed during a given training session. The only difference between the LIWM (Low Interference Working Memory) and HIWM (High Interference Working Memory) tasks concerned the pairs of arms used for each trial in the radial maze during a session. For the LIWM task, different pairs of arms were used for each trial of a given session (4 trials given in 4 different pairs of arms = 8 arms of the radial maze), while for the HIWM task the same pair of arms was used for all trials (equivalent to a Y-maze task). The HIWM task was therefore more repetitive than the LIWM task, with very similar information stored in WM, increasing the level of proactive interference (Underwood, 1957). The third task used in this study was a RM task. During this task, rats had to learn two fixed positions of food rewards in the maze across 8 trials of a given session (equivalent to the 8 runs – 4 trials of 2 phase – in the maze performed by HIWM and LIWM rats during a given session).For the three groups, the behavioral protocol took place on two days, with 10 sessions on Day 1 and 10 sessions on Day 2. For this study, we were interested in the period between these two days of training, during which fEPSPs were continuously recorded, and normalized to the baseline period preceding training. **fEPSP slope analysis.** The initial slope of the fEPSP waveform was analyzed on the baseline and post-training day 1 periods by performing a linear regression on individual fEPSPs between two cursors (previously defined on an average fEPSP) and by computing the coefficient of determination. This analysis was done for each of the valid electrodes among the 8 LFPs electrodes implanted in the DG for an individual rat (see SI Appendix for electrodes selection criteria), and we averaged the results for all these valid electrodes.

### Statistics

Statistical analyses were performed using PRISM *Graphpad Software* version 6. Behavioral data (performance scores expressed as a percentage of correct choices), was analyzed using two-way ANOVAs for repeated measures with Block (2 training sessions) and Group (LIWM, HIWM or RM) as main factors. fEPSP slope was analyzed using two-way ANOVAs for repeated measures with Condition (Baseline or Day 1 post-training) and Time period as main factors. Multiple comparisons were performed using Sidak’s *post hoc* test. Correlations were analyzed using Spearman correlation test due to the non-gaussian distribution of at least one of the variable. Data are expressed as means ± s.e.m. Detailed results of the Statistical tests can be found in SI Appendix, and post-hoc results are represented on the graphs using the standard symbols: * : p ≤ 0.05, ** : p ≤ 0.01, *** : p ≤ 0.001, **** : p ≤ 0.0001.

## Supporting information

Supplemental Materials and methods, figures and statistics

## Acknowledgments

This research was supported by grants from CNRS (ATIP program), *Fondation pour la recherche sur le cerveau* (FRC), and *Région Rhône-Alpes* (CIBLE program). N.F. was also supported by *Fondation pour la recherche médicale* (FRM-FDT20130928087) and *Région Rhône-Alpes* (ARC2 doctoral fellowship). We thank Fernand Malleret for his design and realization of the radial maze apparatus.

## Author Contributions

M.M., N.F., P-A.S. and G.M. designed research; M.M. performed research; M.M., J-C.C., B.T., R.P. and P-A.S. contributed new reagents/analytic tools; M.M. and J-C.C. analyzed data; and M.M. and G.M. wrote the paper.

## References

Awasthi, A., Ramachandran, B., Ahmed, S., Benito, E., Shinoda, Y., Nitzan, N., Heukamp, A., Rannio, S., Martens, H., Barth, J., et al. (2018). Synaptotagmin-3 drives AMPA receptor endocytosis, depression of synapse strength, and forgetting. Science eaav1483.

Barnes, C.A., McNaughton, B.L., Goddard, G.V., Douglas, R.M., and Adamec, R. (1977). Circadian rhythm of synaptic excitability in rat and monkey central nervous system. Science 197, 91–92.

Bear, M.F., and Abraham, W.C. (1996). Long-term depression in hippocampus. Annu. Rev. Neurosci. 19, 437–462.

Bliss, T.V.P., and Lømo, T. (1973). Long-lasting potentiation of synaptic transmission in the dentate area of the anaesthetized rabbit following stimulation of the perforant path. J. Physiol. 232, 331–356.

Bontempi, B., Laurent-Demir, C., Destrade, C., and Jaffard, R. (1999). Time-dependent reorganization of brain circuitry underlying long-term memory storage. Nature 400, 671–675.

Chaillan, F.A., Truchet, B., and Roman, F.S. (2008). Extracellular recordings of rodents in vivo: their contribution to integrative neuroscience. J. Integr. Neurosci. 7, 287–313.

Citri, A., and Malenka, R.C. (2008). Synaptic Plasticity: Multiple Forms, Functions, and Mechanisms. Neuropsychopharmacology 33, 18–41.

Clarke, J.R., Cammarota, M., Gruart, A., Izquierdo, I., and Delgado-García, J.M. (2010). Plastic modifications induced by object recognition memory processing. Proc. Natl. Acad. Sci. U. S. A. 107, 2652–2657.

Cohen, J.S., Reid, S., and Chew, K. (1994). Effects of varying trial distribution, intra- and extramaze cues, and amount of reward on proactive interference in the radial maze. Anim. Learn. Behav. 22, 134–142.

Diekelmann, S., and Born, J. (2010). SLEEP The memory function of sleep. Nat. Rev. Neurosci. 11, 114–126.

Doyère, V., Srebro, B., and Laroche, S. (1997). Heterosynaptic LTD and Depotentiation in the Medial Perforant Path of the Dentate Gyrus in the Freely Moving Rat. J. Neurophysiol. 77, 571–578.

Dudchenko, P.A. (2004). An overview of the tasks used to test working memory in rodents. Neurosci. Biobehav. Rev. 28, 699–709.

Eichenbaum, H., Dudchenko, P., Wood, E., Shapiro, M., and Tanila, H. (1999). The hippocampus, memory, and place cells: is it spatial memory or a memory space? Neuron 23, 209–226.

Fernández-Monreal, M., Brown, T.C., Royo, M., and Esteban, J.A. (2012). The balance between receptor recycling and trafficking toward lysosomes determines synaptic strength during long-term depression. J. Neurosci. Off. J. Soc. Neurosci. 32, 13200–13205.

Fraize, N., Carponcy, J., Joseph, M.A., Comte, J.-C., Luppi, P.-H., Libourel, P.-A., Salin, P.-A., Malleret, G., and Parmentier, R. (2016). Levels of Interference in Long and Short-Term Memory Differentially Modulate Non-REM and REM Sleep. Sleep 39, 2173–2188.

Fraize, N., Hamieh, A.M., Joseph, M.A., Touret, M., Parmentier, R., Salin, P.A., and Malleret, G. (2017). Differential changes in hippocampal CaMKII and GluA1 activity after memory training involving different levels of adaptive forgetting. Learn. Mem. Cold Spring Harb. N 24, 86–94.

Fuster, J.M., and Alexander, G.E. (1971). Neuron activity related to short-term memory. Science 173, 652–654.

Goldman-Rakic, P.S. (1995). Cellular basis of working memory. Neuron 14, 477–485.

Gonzalez, J., Morales, I.S., Villarreal, D.M., and Derrick, B.E. (2014). Low-frequency stimulation induces long-term depression and slow onset long-term potentiation at perforant path-dentate gyrus synapses in vivo. J. Neurophysiol. 111, 1259–1273.

Gruart, A., Muñoz, M.D., and Delgado-García, J.M. (2006). Involvement of the CA3-CA1 synapse in the acquisition of associative learning in behaving mice. J. Neurosci. Off. J. Soc. Neurosci. 26, 1077–1087.

Gruart, A., Sánchez-Campusano, R., Fernández-Guizán, A., and Delgado-García, J.M. (2015). A Differential and Timed Contribution of Identified Hippocampal Synapses to Associative Learning in Mice. Cereb. Cortex N. Y. N 1991 25, 2542–2555.

Hardt, O., Nader, K., and Nadel, L. (2013). Decay happens: the role of active forgetting in memory. Trends Cogn. Sci. 17, 111–120.

Hayashi-Takagi, A., Yagishita, S., Nakamura, M., Shirai, F., Wu, Y.I., Loshbaugh, A.L., Kuhlman, B., Hahn, K.M., and Kasai, H. (2015). Labelling and optical erasure of synaptic memory traces in the motor cortex. Nature 525, 333–338.

Joseph, M.A., Fraize, N., Ansoud-Lerouge, J., Sapin, E., Peyron, C., Arthaud, S., Libourel, P.-A., Parmentier, R., Salin, P.A., and Malleret, G. (2015). Differential Involvement of the Dentate Gyrus in Adaptive Forgetting in the Rat. PloS One 10, e0142065.

Kemp, A., and Manahan-Vaughan, D. (2007). Hippocampal long-term depression: master or minion in declarative memory processes? Trends Neurosci. 30, 111–118.

Loess, H. (1968). Short-term memory and item similarity. J. Verbal Learn. Verbal Behav. 7, 87–92.

Malleret, G., Alarcon, J.M., Martel, G., Takizawa, S., Vronskaya, S., Yin, D., Chen, I.Z., Kandel, E.R., and Shumyatsky, G.P. (2010). Bidirectional regulation of hippocampal long-term synaptic plasticity and its influence on opposing forms of memory. J. Neurosci. Off. J. Soc. Neurosci. 30, 3813–3825.

Matsuo, N., Reijmers, L., and Mayford, M. (2008). Spine-type-specific recruitment of newly synthesized AMPA receptors with learning. Science 319, 1104–1107.

Maviel, T., Durkin, T.P., Menzaghi, F., and Bontempi, B. (2004). Sites of neocortical reorganization critical for remote spatial memory. Science 305, 96–99.

Mi, Y., Katkov, M., and Tsodyks, M. (2017). Synaptic Correlates of Working Memory Capacity. Neuron 93, 323–330.

Migues, P.V., Liu, L., Archbold, G.E.B., Einarsson, E.Ö., Wong, J., Bonasia, K., Ko, S.H., Wang, Y.T., and Hardt, O. (2016). Blocking Synaptic Removal of GluA2-Containing AMPA Receptors Prevents the Natural Forgetting of Long-Term Memories. J. Neurosci. Off. J. Soc. Neurosci. 36, 3481–3494.

Missaire, M., Fraize, N., Joseph, M.A., Hamieh, A.M., Parmentier, R., Marighetto, A., Salin, P.A., and Malleret, G. (2017). Long-term effects of interference on short-term memory performance in the rat. PloS One 12, e0173834.

Miyashita, Y. (1988). Neuronal correlate of visual associative long-term memory in the primate temporal cortex. Nature 335, 817–820.

Mongillo, G., Barak, O., and Tsodyks, M. (2008). Synaptic theory of working memory. Science 319, 1543–1546.

Moser, E.I., Moser, M.B., and Andersen, P. (1994). Potentiation of dentate synapses initiated by exploratory learning in rats: dissociation from brain temperature, motor activity, and arousal. Learn. Mem. Cold Spring Harb. N 1, 55–73.

Nabavi, S., Fox, R., Proulx, C.D., Lin, J.Y., Tsien, R.Y., and Malinow, R. (2014). Engineering a memory with LTD and LTP. Nature 511, 348–352.

Nanry, K.P., Mundy, W.R., and Tilson, H.A. (1989). Colchicine-induced alterations of reference memory in rats: role of spatial versus non-spatial task components. Behav. Brain Res. 35, 45–53.

Nicholls, R.E., Alarcon, J.M., Malleret, G., Carroll, R.C., Grody, M., Vronskaya, S., and Kandel, E.R. (2008). Transgenic mice lacking NMDAR-dependent LTD exhibit deficits in behavioral flexibility. Neuron 58, 104–117.

Niewoehner, B., Single, F.N., Hvalby, Ø., Jensen, V., Meyer zum Alten Borgloh, S., Seeburg, P.H., Rawlins, J.N.P., Sprengel, R., and Bannerman, D.M. (2007). Impaired spatial working memory but spared spatial reference memory following functional loss of NMDA receptors in the dentate gyrus. Eur. J. Neurosci. 25, 837–846.

O’Keefe, J. (1993). Hippocampus, theta, and spatial memory. Curr. Opin. Neurobiol. 3, 917–924.

Park, P., Volianskis, A., Sanderson, T.M., Bortolotto, Z.A., Jane, D.E., Zhuo, M., Kaang, B.-K., and Collingridge, G.L. (2014). NMDA receptor-dependent long-term potentiation comprises a family of temporally overlapping forms of synaptic plasticity that are induced by different patterns of stimulation. Philos. Trans. R. Soc. B-Biol. Sci. 369, 20130131.

Pavlowsky, A., Wallace, E., Fenton, A.A., and Alarcon, J.M. (2017). Persistent modifications of hippocampal synaptic function during remote spatial memory. Neurobiol. Learn. Mem. 138, 182–197.

Poirier, G.L., Amin, E., and Aggleton, J.P. (2008). Qualitatively different hippocampal subfield engagement emerges with mastery of a spatial memory task by rats. J. Neurosci. Off. J. Soc. Neurosci. 28, 1034–1045.

Rainer, G., and Miller, E.K. (2002). Timecourse of object-related neural activity in the primate prefrontal cortex during a short-term memory task. Eur. J. Neurosci. 15, 1244–1254.

Rasch, B., and Born, J. (2013). About sleep’s role in memory. Physiol. Rev. 93, 681–766.

Roberts, W.A., and Dale, R.H.I. (1981). Remembrance of places lasts: Proactive inhibition and patterns of choice in rat spatial memory. Learn. Motiv. 12, 261–281.

Ryan, T.J., Roy, D.S., Pignatelli, M., Arons, A., and Tonegawa, S. (2015). Engram cells retain memory under retrograde amnesia. Science 348, 1007–1013.

Sasaki, T., Piatti, V.C., Hwaun, E., Ahmadi, S., Lisman, J.E., Leutgeb, S., and Leutgeb, J.K. (2018). Dentate network activity is necessary for spatial working memory by supporting CA3 sharp-wave ripple generation and prospective firing of CA3 neurons. Nat. Neurosci. 21, 258–269.

Shafi, M., Zhou, Y., Quintana, J., Chow, C., Fuster, J., and Bodner, M. (2007). Variability in neuronal activity in primate cortex during working memory tasks. Neuroscience 146, 1082–1108.

Smith, C. (1996). Sleep states, memory processes and synaptic plasticity. Behav. Brain Res. 78, 49–56.

Takeuchi, T., Duszkiewicz, A.J., and Morris, R.G.M. (2014). The synaptic plasticity and memory hypothesis: encoding, storage and persistence. Philos. Trans. R. Soc. B-Biol. Sci. 369, 20130288.

Tanaka, K.Z., He, H., Tomar, A., Niisato, K., Huang, A.J.Y., and McHugh, T.J. (2018). The hippocampal engram maps experience but not place. Science 361, 392–397.

Tolman, E.C. (1925). Purpose and cognition: the determiners of animal learning. Psychol. Rev. 32, 285–297.

Tonegawa, S., Liu, X., Ramirez, S., and Redondo, R. (2015). Memory Engram Cells Have Come of Age. Neuron 87, 918–931.

Tononi, G., and Cirelli, C. (2014). Sleep and the Price of Plasticity: From Synaptic and Cellular Homeostasis to Memory Consolidation and Integration. Neuron 81, 12–34.

Underwood, B.J. (1957). Interference and forgetting. Psychol. Rev. 64, 49–60.

Whitlock, J.R., Heynen, A.J., Shuler, M.G., and Bear, M.F. (2006). Learning induces long-term potentiation in the hippocampus. Science 313, 1093–1097.

Wixted, J.T. (2004). The psychology and neuroscience of forgetting. Annu. Rev. Psychol. 55, 235–269.

Xavier, G.F., and Costa, V.C.I. (2009). Dentate gyrus and spatial behaviour. Prog. Neuropsychopharmacol. Biol. Psychiatry 33, 762–773.

Xavier, G.F., Oliveira-Filho, F.J., and Santos, A.M. (1999). Dentate gyrus-selective colchicine lesion and disruption of performance in spatial tasks: difficulties in “place strategy” because of a lack of flexibility in the use of environmental cues? Hippocampus 9, 668–681.

