## Supplemental Materials and methods, figures and statistics for "Working and Reference Memory tasks trigger opposed long-term synaptic changes in the rat dentate gyrus"

**This PDF file includes:**

Main Text
Figures 1 to 2
Tables 1 to 4

### Supplemental Materials and Methods

**Animals.** 33 Dark Agouti male rats (*Janvier Labs*) aged 10 weeks at the time of surgery (weigh of about 200g) were used in this study. Animals were housed individually after surgery in a ventilated cabinet with a 12h/12h (9am-9pm) light/dark cycle at 24°C and *ad libitum* access to food (except during food deprivation periods necessary for the behavioral tasks) and water. This study was carried out in strict accordance with the recommendations of the Lyon 1 University (CE2A-UCBL 55) and the European (2010/63/UE) ethical committees for the use of experimental animals. The protocol was approved by the Lyon 1 University ethical committee (Permit Number: DR2016-29). All efforts were made to minimize the number of animals used and their suffering during the experimental procedures.

**Electrodes.** All the electrodes used in this article were made on site. The two EEG electrodes and the reference electrode were made of a stainless steel wire (76µm-diameter) (*PHYMEP*) welded to a stainless steel screw. EMG electrodes were also made of the 76µm diameter stainless steel wire, at the end of which was formed a gold-plated tin ball (around 1mm-diameter). Each of the 8 LFP electrodes was made of a single non-bared tungsten wire (45µm-diameter, *California Fine Wire*, U.S.A.), and were arranged in a recording array on two rows of 4 electrodes (100µm between two adjacent electrodes) with a final dimension of around 300 x 700 µm. The bipolar stimulating electrode was made of two twisted stainless steel wires (100µm-diameter, *California Fine Wire*, U.S.A.), parted on the last 400µm and bared on 200µm.

**Stereotaxic surgery.** Initial anesthesia of the rats was performed in an induction chamber saturated with isoflurane (2-2.5%). Rats were then placed on the stereotaxic frame and anesthesia was maintained with a gas mix containing 0.75-1% of isoflurane and enriched in oxygen. After scalp incision, 5 craniotomies were performed at the position of the different electrodes, with EEG on the left side of the skull (1 prefrontal EEG + 1 parietal EEG) and LFP and stimulating electrodes on the right side on the skull. Two EMG electrodes were inserted between the neck muscles of the rat. The PP (perforant path) stimulating electrode was lowered at -7.5mm (Antero-Posterior) and +4mm (Medio-lateral) relative to the Bregma with a slow descending rate of 0.05mm per minute until -3mm (Dorso-Ventral). The recording array of 8 LFP electrodes was lowered simultaneously at -3.3mm (AP) and +2.4mm (ML) relative to Bregma, and the electrophysiological signals were recorded during the slow descent (0.05mm per minute). During this descent, signals from CA1 *stratum pyramidale*were clearly recognizable from a high spiking activity at a depth of around -2.5mm. Electrical stimulations of the PP started when both the stimulating electrode and the recording array were around -3mm (DV), and consisted in a monophasic 200µs pulse delivered every 3 seconds (stimulation intensity around 90µA). The stimulating electrode and the recording array were then lowered concomitantly and slowly until we could visualize a stereotyped fEPSP from the PP-DG synapse with a delay to the first peak inferior to 5ms, a positive polarity and a maximized amplitude (average depth of the recording array and the stimulating electrode = -3.5mm). Once the optimal depths of the electrodes were reached, stimulations went on for at least 20 minutes in order to ensure the stable position of the electrodes, which were then fixed to the skull using Super-Bond (*C&B*), and linked to a connector (*EIB-27, Neuralynx, U.S.A.*) along the EEG and EMG electrodes. The connector was protected and cemented to the skull using dental cement, and skin was sutured around this protective “hat”. At the end of surgery, rats were subcutaneously injected with 3mL of glucose (2.5%) containing carprofen (5mg/kg). Rats were then allowed to recover during 10-15 days in individual cages, and were daily weighed and habituated to be manipulated by the experimenter.

**Recording setup.** During the experimental protocol, rats were recorded individually in a mobile opaque cage (57x39x50 cm) placed on the radial maze and with no cover to allow the mobility of the recording cable fixed on the ceiling of the room (as rats stayed plugged to the cable for the whole duration of the protocol, including while exploring the radial maze). The light ON-OFF cycle and the temperature in this recording room was the same as in the housing ventilated cabinet (12/12h, 9am to 9pm, 24°C). Before the beginning of the protocol, rats were habituated during several days to the recording setup. Signals were acquired through 1) a homemade preamplifier plugged on the EIB connector, 2) a recording cable plugged to a rotating connector (*Plastic One Inc. CT*), 3) a homemade connector referencing all LFPs and EEGs signals to the reference electrode, 4) a 16-Channel Amplifier (*AM-Systems, U.S.A.*) allowing amplification (x1000) and filtration (0.3 - 1000 Hz for LFPs) of the signals, 5) a data acquisition card (*NI-6343, National Instruments, U.S.A*.) wih a 5000Hz sampling frequency, 6) a computer with a MATLAB Software (*The MathWorks, U.S.A*.) storing data for offline analysis.

**Behavioral apparatus and habituation.** After 10-15 days of recovery post-surgery, rats were food deprived for a week in order to reach 80% of their baseline weight. During this period and during the following maze habituation period, rats were habituated to the recording setup by being recorded overnight in the recording cage. During the maze habituation period of 8 days, rats were trained in 3-4 sessions everyday (between 9am and 12pm) to collect food rewards in all 8 arms of the radial maze, first without the recording cable, and then while being plugged to the cable. The radial maze was made of 8 arms (65cm x 12cm) around a central octagonal platform (33cm diameter). A rectangular platform (17cm x 25 cm) was placed at the end of each arm, and the arms could be automatically risen (allowing access to the platforms) or lowered (preventing access to the platforms) by an experimenter who could follow the rat's movements in the maze from an adjacent room using a camera above the maze. On each of the 8 peripheral platforms was placed a cube (2x2cm and 0.5cm-deep) used as a food well for food pellets (*Dustless Precision Pellets;Bioserve, Frenchtwon, NJ*) odorless and invisible for the rat from the central platform. The behavioral room contained many distal visual cues (furniture, door... etc) that could be used by rats to navigate in the maze using allocentric spatial hippocampal-dependent memory (Eichenbaum, 1999; O'Keefe, 1993; Poirier et al., 2008). Once rats were well-habituated to collect food rewards in the maze (<2 minutes to visit the 8 arms), the experimental protocol could begin.

**Outline of the experimental protocol.** Rats were first re-habituated for 24h to the recording setup. The next day, an input-output (I/O) curve was established to choose the stimulation intensity for the whole protocol. Stimulations were then started (1 stimulation/30 seconds) and fEPSPs could thus stabilize (their amplitude was checked regularly) until the start of the baseline recordings on the next day. We fixed the start of the Baseline recordings the following day at 9am (ZT0, when lights were turned on in the recording room), and checked the fEPSPs regularly to make sure that they were stable for 24h. When it was the case, we could start the first training day (in one of the three behavioral tasks) that took place between 9am (ZT0) and 12pm (ZT3), during which stimulations were stopped. After training, rats were allowed to rest in the recording cage, and stimulations resumed every 30 seconds for 21h until the following day when the second training day was performed between 9am and 12pm (stimulations stopped).

**Behavioral protocols.** The sample and choice phase of the LIWM and HIWM tasks are described in the main manuscript. For each phase, the rat started from one of the two arms opposed (by central symmetry) to the pair of arms used for the choice phase. This start from an arm and not from the central platform allowed rats to enter the central platform facing the pair of arms from which they had to choose to enter only one of them (avoiding impulsive decision-making). Moreover, the starting arm was chosen pseudo-randomly for each phase and each trial, in order to avoid the used of motor strategies (turn right, turn left, go straight). The starting arm could thus be the same or be different for the two phases (sample or choice phase) of the same trial. Moreover, the order of presentation of the sample and choice arms was also pseudo-randomized so that half of the trials of a given session involve a correct choice when choosing the right arm of the pair (and the other half for the left arm of the pair). The behavioral score was comprised between 0 and 4 for these two WM tasks (4 trials of 2 phases per *session* = 8 runs per *session*).

We also used pseudo-randomly chosen starting arms in the RM task, from which rats could start during one trial(except the two rewarded arms). Moreover, we prevented the occurrence of WM errors in this RM task by lowering all previously visited arms during an ongoing session at the beginning of each trial, until the two rewarded arms were visited. When the two rewards were collected, we refilled the food pellets, opened again all arms of the maze, and proceeded as before until the end of the session. The behavioral score was comprised between 2 and 8 (8 trials with a single choice phase, with at least one visit in each of the two rewarded arms). Contrary to the LIWM and HIWM tasks during which rats always collected at least 4 rewards during one session due to the 4 sample phases, RM rats could sometimes collect only two rewards in one session (especially at the beginning of training because this task is more complex than the WM tasks, and the behavioral score was generally comprised between 2 and 4 in the first sessions). We therefore balanced the number of rewards collected during one session between the three tasks by increasing the number of food pellets in the food wells from 1 (as in the LIWM and HIWM tasks) to 2-3 in the RM tasks at the beginning of training. This number of food pellets was then decreased as rats learned the task.

For the three tasks, a 15-second delay separated two runs, during which rats were placed in the opaque transfer cage placed on the central platform of the maze. Similarly, rats were placed in this transfer cage during the 10 minutes between two training sessions, and were kept awake during this delay.

**Dataset, histological verification and exclusions.** Rats were included in this study only after histological verification of electrode position. To do so, an electrocoagulation procedure was performed on the rats before their sacrifice with intraperitoneal injection of Dolethal. Briefly, rats were anesthetized with 2% isoflurane, and an electrical stimulation was applied between each LFP electrode and the reference electrode (1s, 600µA), and between the two poles of the stimulation electrode (0.5s, 500µA). The brains of the rats were then frozen at -40°C, and transverse sections (40µm) were obtained and colored with neutral red staining. Correct electrode placement (in the DG and PP) was then verified using the atlas of Paxinos and Watson (2007).

In addition to this histologic verification, an electrophysiological verification was realized in order to select the correct electrodes among the 8 implanted LFP electrodes in each rat. Only the electrodes displaying stereotyped PP-DG fEPSPs (positive polarity, delay to the first peak inferior to 5.3ms and a classic waveform with the presence of only one population spike more or less visible) were selected, for both fEPSP and spontaneous oscillations analyses. The number of selected electrodes *per* rat varied between 2 and 8 (mean 6.7 ± 1.4 standard deviation). To prevent biases by selecting a given electrode for a given rat, we averaged all results (fEPSP, spontaneous oscillations) on all the selected correct electrodes for each rat.

Concerning the fEPSP analysis, we excluded from the data set individual fEPSPs that displayed 1) a $r^2$ inferior to 0.5 reflecting potential artefactual responses, 2) a slope below or above the mean fEPSP slope on 24h ± 3 standard deviation.

### Supplemental Figures

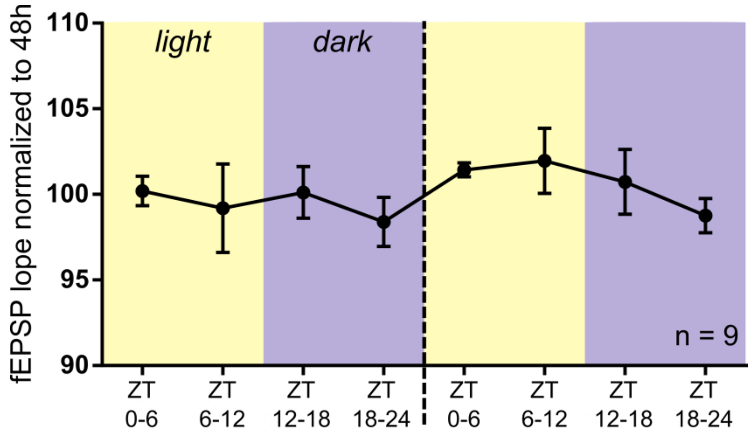


**Fig.S1.** Circadian modulation of the fEPSP slope at PP-DG synapse. Stable baseline fEPSPs were recorded for 48h from 9 rats before training. This graph displays the evolution of the fEPSP slope during these 48 hours. Note the tendency for a decrease in the fEPSP slope during a given day (ZT0-6 to ZT18-24), followed by an increase of the fEPSP slope between the end of the first day and the beginning of the second day when light is turned on. Data were not normal (D'agostino & Peason omnibus normality test: K2 = 16.47; p = 0.0003), so we used the Friedman test which only showed a tendency for an effect of the time periods (Friedman statistic: 12.04; p = 0.0993) due to the low number of animals. Data are represented as mean ± SEM.

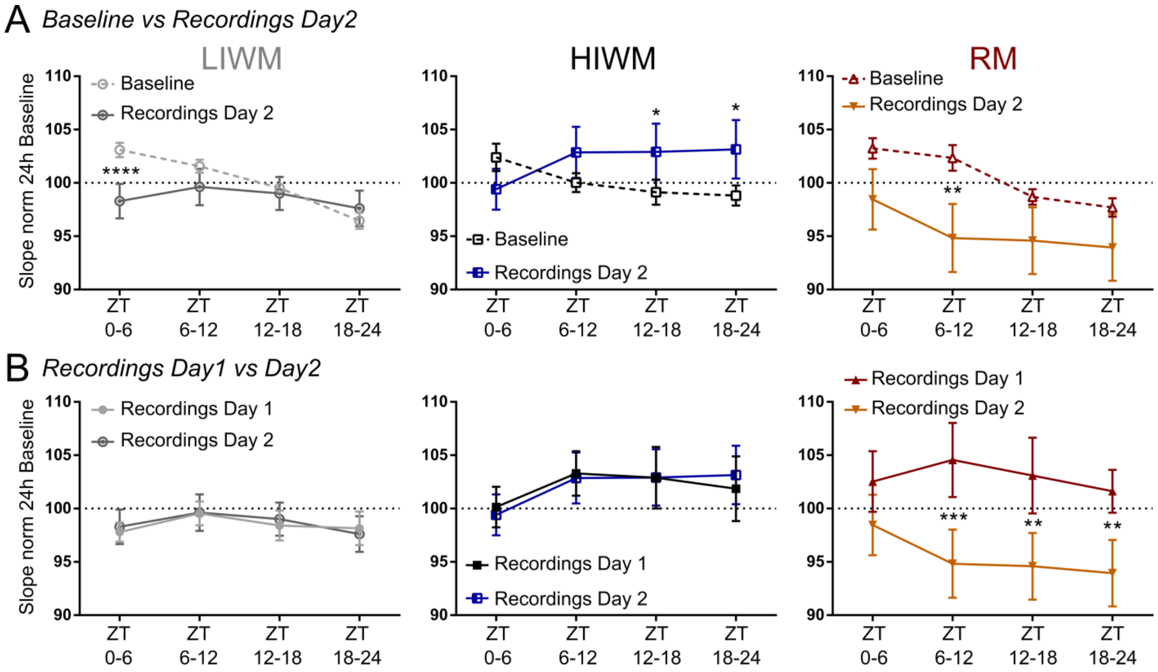


**Fig.S2.** Synaptic changes recorded on Day 2 after training in the three tasks, compared to Baseline and to Day 1 recordings. Direct within-period comparison of the fEPSP slope (normalized to the averaged slope on 24h of Baseline) between Baseline and Day 2 recordings *(A)*, or between Day 1 and Day 2 recordings *(B)*. For the LIWM group, the repeated-measure ANOVA 2 and post-hoc tests revealed that on Day 2 post-training, fEPSP slope at PP-DG synapse remains stable and at the same level as on Day 1 *(B)*, with therefore the same significant early synaptic depression compared to Baseline at ZT3-6 (first 3 hours post-training) *(A)*. Similarly for the HIWM group, synaptic transmission on Day 2 post-training was stable and at the same level as on Day 1 *(B)*, and we find again a significant late synaptic potentiation compared to Baseline, starting 9h and ending 21h post-training (ZT12-18 – ZT18-24) *(A)*. Interestingly, for the RM group, we found opposed long-term synaptic changes after training Day 1 and 2. Indeed, we previously found a significant late synaptic potentiation after the first day of RM training (Figure 3), which is followed here by a synaptic depression after the second day of training. Indeed, the repeated-measure ANOVA 2 and post-hoc tests revealed a slight synaptic depression between Baseline and Day 2 recordings, especially at ZT6-12 *(A)*, whereas this synaptic depression is even more pronounced between Day 1 and Day 2 recordings, starting 3h post-training and lasting for the whole day until 21h post-training *(B)*. Data are represented as mean ± SEM.

### Statistical results

RpM(...) = Repeated Measure (factor concerned by repetition)

**Table 1.** Statistical results of Figure 2.

| Graph | Statistical test | Factor | Statistics | p-value |
|---|---|---|---|---|
| (A) | RpM (Blocks) ANOVA 2 Day 1 | Block | $F_{4,176} = 23.84$ | < 0.0001 **** |
| | | Group | $F_{2,44} = 35.14$ | < 0.0001 **** |
| | | Block x Group | $F_{8,176} = 2.674$ | 0.0085 ** |
| | RpM (Blocks) ANOVA 2 Day 2 | Block | $F_{4,172} = 4.302$ | < 0.0024 ** |
| | | Group | $F_{2,43} = 65.17$ | < 0.0001 **** |
| | | Block x Group | $F_{8,172} = 4.721$ | < 0.0001 **** |
| (B) | RpM (Days) ANOVA 2 | Day | $F_{1,43} = 156.8$ | < 0.0001 **** |
| | | Group | $F_{2,43} = 25.91$ | < 0.0001 **** |
| | | Day x Group | $F_{2,43} = 67.74$ | < 0.0001 **** |
| (C) | RpM (Trials) ANOVA 2 S1-5 | Trial | $F_{1,28} = 0.1550$ | 0.6968 |
| | | Group | $F_{1,28} = 1.383$ | 0.2495 |
| | | Trial x Group | $F_{1,28} = 1.792$ | 0.1915 |
| | RpM (Trials) ANOVA 2 S6-10 | Trial | $F_{1,28} = 1.412$ | 0.2448 |
| | | Group | $F_{1,28} = 13.24$ | 0.0011 ** |
| | | Trial x Group | $F_{1,28} = 1.412$ | 0.2448 |
| | RpM (Trials) ANOVA 2 S11-15 | Trial | $F_{1,27} = 6.818$ | 0.0146 * |
| | | Group | $F_{1,27} = 25.21$ | < 0.0001 **** |
| | | Trial x Group | $F_{1,27} = 1.182$ | 0.2866 |
| | RpM (Trials) ANOVA 2 S16-20 | Trial | $F_{1,27} = 0.05976$ | 0.8087 |
| | | Group | $F_{1,27} = 112.1$ | < 0.0001 **** |
| | | Trial x Group | $F_{1,27} = 0.2165$ | 0.6455 |

**Table 2.** Statistical results of Figure 3.

| Graph | Statistical test | Factor | Statistics | p-value |
|---|---|---|---|---|
| (A) | RpM (Time) ANOVA 2 | Time | $F_{3,108} = 15.45$ | < 0.0001 **** |
| | | Group | $F_{2,36} = 2.015$ | 0.1481 |

| | | | | |
|---|---|---|---|---|
| | Baseline | Time x Group | $F_{6,108} = 0.9199$ | 0.4837 |
| | RpM (Time) ANOVA 2 Day 1 | Time | $F_{3,108} = 1.944$ | 0.1268 |
| | | Group | $F_{2,36} = 1.213$ | 0.3091 |
| | | Time x Group | $F_{6,108} = 0.2668$ | 0.9512 |
| (B) | RpM (both) ANOVA 2 LIWM | Time | $F_{3,36} = 6.177$ | 0.0017 ** |
| | | Condition | $F_{1,12} = 2.550$ | 0.1363 |
| | | Time x Condition | $F_{3,36} = 9.490$ | < 0.0001 **** |
| | RpM (both) ANOVA 2 HIWM | Time | $F_{3,33} = 0.1692$ | 0.9164 |
| | | Condition | $F_{1,11} = 0.8849$ | 0.3671 |
| | | Time x Condition | $F_{3,33} = 5.533$ | 0.0034 ** |
| | RpM (both) ANOVA 2 RM | Time | $F_{3,39} = 3.910$ | 0.0156 * |
| | | Condition | $F_{1,13} = 0.8795$ | 0.3654 |
| | | Time x Condition | $F_{3,39} = 2.439$ | 0.0789 |
| (C) | RpM (Time) ANOVA 2 Baseline | Time | $F_{3,108} = 13.05$ | < 0.0001 **** |
| | | Group | $F_{2,36} = 0.03859$ | 0.9622 |
| | | Time x Group | $F_{6,108} = 1.950$ | 0.0792 |
| | RpM (Time) ANOVA 2 Day 1 | Time | $F_{3,108} = 3.396$ | 0.0205 * |
| | | Group | $F_{2,36} = 0.9178$ | 0.4085 |
| | | Time x Group | $F_{6,108} = 0.5326$ | 0.7824 |
| (D) | RpM (both) ANOVA 2 LIWM | Time | $F_{3,36} = 3.004$ | 0.0430 * |
| | | Condition | $F_{1,12} = 26.37$ | 0.0002 *** |
| | | Time x Condition | $F_{3,36} = 2.802$ | 0.0537 |
| | RpM (both) ANOVA 2 HIWM | Time | $F_{3,33} = 1.855$ | 0.1564 |
| | | Condition | $F_{1,11} = 7.942$ | 0.0167 * |
| | | Time x Condition | $F_{3,33} = 1.635$ | 0.2001 |
| | RpM (both) ANOVA 2 RM | Time | $F_{3,39} = 1.201$ | 0.3222 |
| | | Condition | $F_{1,13} = 12.18$ | 0.0040 ** |
| | | Time x Condition | $F_{3,39} = 5.122$ | 0.0044 ** |

**Table 3.** Statistical results of Figure 4.

| Graph | Group | Time period | Spearman's rho | p-value |
|---|---|---|---|---|
| (A) | LIWM | ZT3-6 | -0.3344 | 0.0186 * |
| | | ZT6-12 | 0.2006 | 0.5163 |

| | | | | |
|---|---|---|---|---|
| | | ZT12-18 | 0.08174 | 0.4580 |
| | | ZT18-24 | 0.3195 | 0.2902 |
| | HIWM | ZT3-6 | -0.2714 | 0.2661 |
| | | ZT6-12 | -0.1923 | 0.3962 |
| | | ZT12-18 | -0.5731 | 0.0264 * |
| | | ZT18-24 | -0.4750 | 0.0669 |
| | RM | ZT3-6 | 0.1581 | 0.5864 |
| | | ZT6-12 | 0.1181 | 0.6861 |
| | | ZT12-18 | -0.1403 | 0.5974 |
| | | ZT18-24 | -0.1314 | 0.6192 |
| (B) | LIWM | ZT3-6 | -0.1480 | 0.5422 |
| | | ZT6-12 | -0.3018 | 0.2559 |
| | | ZT12-18 | -0.4042 | 0.1322 |
| | | ZT18-24 | -0.09964 | 0.6544 |
| | HIWM | ZT3-6 | -0.1120 | 0.6516 |
| | | ZT6-12 | -0.04696 | 0.8096 |
| | | ZT12-18 | -0.6718 | 0.0128 * |
| | | ZT18-24 | -0.5382 | 0.0562 |
| | RM | ZT3-6 | -0.01333 | 0.9360 |
| | | ZT6-12 | 0.09556 | 0.7444 |
| | | ZT12-18 | 0.2511 | 0.3836 |
| | | ZT18-24 | 0.5023 | 0.0693 |
| (C) | LIWM | ZT3-6 | 0.2400 | 0.4478 |
| | | ZT6-12 | 0.1904 | 0.5494 |
| | | ZT12-18 | 0.1409 | 0.6591 |
| | | ZT18-24 | -0.2895 | 0.2266 |
| | HIWM | ZT3-6 | -0.2004 | 0.4787 |
| | | ZT6-12 | -0.07158 | 0.7652 |
| | | ZT12-18 | 0.2290 | 0.4712 |
| | | ZT18-24 | 0.1825 | 0.5681 |
| | RM | ZT3-6 | 0.2843 | 0.3207 |
| | | ZT6-12 | 0.5875 | 0.0301 * |
| | | ZT12-18 | 0.5616 | 0.0396 * |

| | | | | |
|---|---|---|---|---|
| | | ZT18-24 | 0.5851 | 0.0309 * |

**Table 4.** Statistical results of Figure S2.

| Graph | Statistical test | Factor | Statistics | p-value |
|---|---|---|---|---|
| (A) | RpM (both) ANOVA 2 LIWM | Time | $F_{3,36}$ = 13.30 | < 0.0001 **** |
| | | Condition | $F_{1,12}$ = 0.9762 | 0.3426 |
| | | Time x Condition | $F_{3,36}$ = 7.702 | 0.0004 *** |
| | RpM (both) ANOVA 2 HIWM | Time | $F_{3,33}$ = 0.06607 | 0.9775 |
| | | Condition | $F_{1,11}$ = 0.7716 | 0.3985 |
| | | Time x Condition | $F_{3,33}$ = 6.351 | 0.0016 ** |
| | RpM (both) ANOVA 2 RM | Time | $F_{3,39}$ = 3.382 | 0.0276 * |
| | | Condition | $F_{1,13}$ = 4.259 | 0.0596 |
| | | Time x Condition | $F_{3,39}$ = 0.6275 | 0.6016 |
| (B) | RpM (both) ANOVA 2 LIWM | Time | $F_{3,36}$ = 1.972 | 0.1356 |
| | | Condition | $F_{1,12}$ = 0.01712 | 0.8981 |
| | | Time x Condition | $F_{3,36}$ = 0.3000 | 0.8252 |
| | RpM (both) ANOVA 2 HIWM | Time | $F_{3,33}$ = 1.777 | 0.1706 |
| | | Condition | $F_{1,11}$ = 0.000423 | 0.9840 |
| | | Time x Condition | $F_{3,33}$ = 0.3021 | 0.8237 |
| | RpM (both) ANOVA 2 RM | Time | $F_{3,39}$ = 0.7898 | 0.5069 |
| | | Condition | $F_{1,13}$ = 5.239 | 0.0395 * |
| | | Time x Condition | $F_{3,39}$ = 1.298 | 0.2887 |

**References**


Eichenbaum, H. (1999). The hippocampus and mechanisms of declarative memory. Behav. Brain Res. *103*, 123–133.

O'Keefe, J. (1993). Hippocampus, theta, and spatial memory. Curr. Opin. Neurobiol. *3*, 917–924.

Poirier, G.L., Amin, E., and Aggleton, J.P. (2008). Qualitatively different hippocampal subfield engagement emerges with mastery of a spatial memory task by rats. J. Neurosci. Off. J. Soc. Neurosci. *28*, 1034–1045.
